# Fast and accurate short read alignment with hybrid hash-tree data structure

**DOI:** 10.1101/2024.02.20.581311

**Authors:** Junichiro Makino, Toshikazu Ebisuzaki, Ryutaro Himeno, Yoshihide Hayashizaki

**Affiliations:** Advanced Accelerating Systems Co. Ltd., Deiki 1-28, B1312, Kanazawa-ku, Yokohama, Kanagawa 236-0021 Japan; Department of Planetology, Graduate School of Science, Kobe University, 1-1, Rokkodai-cho, Nada-ku, Kobe 657-8051 Japan; K.K. Dnaform, Ask Sanshin Building 3F, 2-6-29 Tsurumi-chuo, Tsurumi-ku, Tsurumi, Yokohama, Kanagawa, 230-0051 Japan; Juntendo University, Faculty of Health Data Science, 6-8-1 Hinode, Urayasu, Chiba 279-0013 Japan; Juntendo University, Medical Technology Innovation Center, 2-1-1 Hongo, Bunkyo-ku, Tokyo 113-8421 Japan

**Keywords:** human whole genome analysis, short read, alignment (mapping), variant calling, hash, tree

## Abstract

Rapidly increasing amount of short read data generated by NGSs (new-generation sequencers) calls for the development of fast and accurate read alignment programs. The programs based on hash table (BLAST) and Burrows-Wheeler transform (bwa-mem) are used, and the latter is known to give superior performance. We here present a new algorithm, a hybrid of hash table and suffix tree, which we designed to speed up the alignment of short reads against large reference sequences such as human genome. The total turnaround time for processing one human genome sample (read depth of 30) is just 31 minutes with our system while that was more than 25 hours with bwa-mem/gatk. The time for aligner alone is 28 minutes for our system but around 2 hours for bwa-mem. Our new algorithm is 4.4 times faster than bwa-mem while achieving similar accuracy. Variant calling and other downstream analyses after the alignment can be done with open-source tools such as SAMtools and Genome Analysis Toolkit (gatk) packages, as well as our own fast variant caller, which is well parallelized and much faster than gatk.

## Introduction

Present-day sequencers such as Illumina NextSeq and BGI T7 can produce full read of human genome of around 50 persons (read depth of 30) in one day. This data corresponds to around 10TBp (base pair). This enormous amount of data requires a new level of computational power for read alignment and variant calling. The current “best practice” pipeline uses bwa-mem (Li, 2012; 2013) for alignment and gatk (Poplin et al., 2018) for variant calling. The use of these tools on usual CPU-based servers would require a large cluster system to handle the output of a single sequencer, resulting in the significant increase of the total cost which would compromise the advantage of modern sequencers. Thus, it is of critical importance to improve the performance of read alignment and variant calling either by improving the hardware, software or both.

There are many works to improve the performance of human genome analysis, mostly by using faster processors. Here we discuss a few recent achievements. A proprietary implementation of these tools on a machine with four NVIDIA V100 GPUs realized the speed of 175 minutes for a human genome data of read depth of 50 (Franke and Crowgey, 2020). This speed corresponds to 1.2 TBp/day. Thus, we can construct a system which has the performance of 10 TBp/day using 4 × 8 = 32 GPUs (or a somewhat smaller number with newer GPUs such as A100 or H100). Such a system would still be pretty expensive and would consume a large amount of electricity. This implementation is not a simple porting of bwa-mem and gatk but newly developed code highly optimized to NVIDIA GPUs. Therefore, the results of their system are not exactly the same as those of the best practice pipeline. As a result, they gave detailed discussions on the accuracy of their system.

An implementation of bwa-mem and gatk on Supercomputer Fugaku with Fujitsu A64fx processor has been reported in Suzuki et al. (2021). The achieved performance is around 200Gbps/h, or 5TBp/day using 96 nodes of Supercomputer Fugaku. Thus, in principle a 192-node A64fx system can process the data from one sequencer, but such a system is quite expensive and requires too much space and electricity. This implementation is a straightforward porting of bwa-mem and gatk to the A64fx processor of Supercomputer Fugaku with some modification of the source code to make use of the SVE SIMD instruction set of the A64fx processor. Thus, the calculation results are the same as those of the original bwa-mem/gatk combination, and there is no need for detailed accuracy comparison. Since the gatk is not well parallelized, they have implemented a fairly complex scheduling algorithm in which multiple samples are processed in one batch, so that they could improve the parallel efficiency.

Illumina provides the hardware-based acceleration system, Dragen, which realizes the throughput of around 10TBp/day. This system apparently offers the best price-performance ratio for human genome analysis. They offer both the software-only and hardware-accelerated versions of their systems.

In summary, it is certainly possible to construct computer systems which can process the data from a modern sequencer real time, but such systems are very expensive. To fully utilize the high performance of modern sequencers for the analysis of human genome, it is necessary to significantly improve the performance of both the alignment and variant calling. The latter can be achieved by implementing the basic valiant calling algorithms in efficient parallel programs, while the former requires a fundamentally new algorithm, if we are to achieve such improvement over existing best-performing software.

We describe such a new algorithm, the hybrid hash-tree algorithm. In this paper, we describe this new algorithm and compare its performance and accuracy with those of bwa-mem/gatk combination. The new algorithm achieved much better performance, while retaining the accuracy comparable to that of bwa-mem/gatk.

## Methods

### Hash-based algorithm

The hybrid hash-tree algorithm is based on the original hash-based algorithm such as used in BLAST (Altschul et al., 1990; Camacho et al., 2009). A hash key is a fixed-length sequence of bases. With the hash-based algorithm, for each of all possible hash keys of a given length *l*, the locations in the reference sequence which match that key are recorded. For a read, we first use its first *l* bases as a key. We store the locations of this key in the reference. Then, we shift the starting position within the read by stride *s* (typically *s*= 5), and store the locations the new key. We repeat this procedure until we reach the end of the read.

Then we sort all candidate locations, and search for the locations at which several different positions in the read match to the reference sequence. For example, if the read is a perfect copy of one sub-sequence of the reference sequence and if this sub-sequence appears no other location, each hash of the read would appear the corresponding location of the reference sub-sequence and there is no other place in the reference genome where all hashes of the read appear the corresponding location of the sub-sequence. Thus, we can determine the location of the sequence.

Let us consider the reference sequence of AGTCACCAGAGATGGC with length 16 as an example. With *l* = 2, we have 15 possible starting locations of keys. If we encode ACGT as 0, 1, 2, 3, the locations and keys are as shown in figure 1.

**Figure 1.**
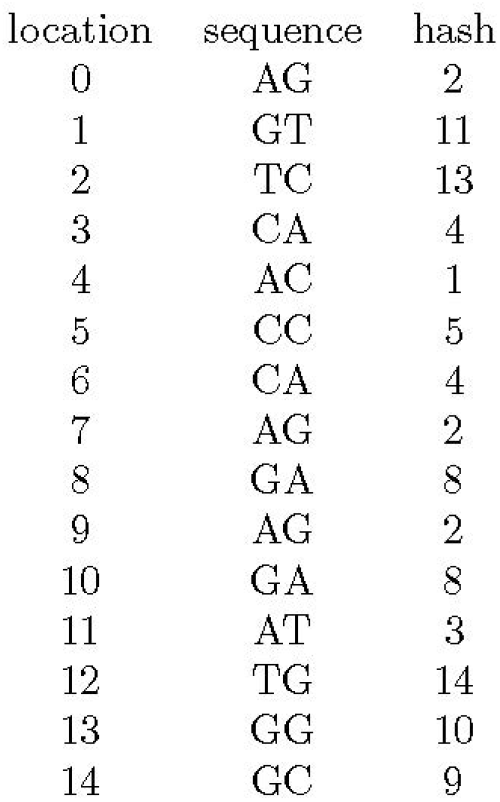
Hash key values for 15 locations of reference sequence AGTCACCAGAGATGGC. First, second and third columns show the location, key value, and original string.

Therefore, the hash table should express the data structure shown in figure 2. If the key is, for example, 0, it should return NULL, since there is no sequence AA in the reference.

**Figure 2.**
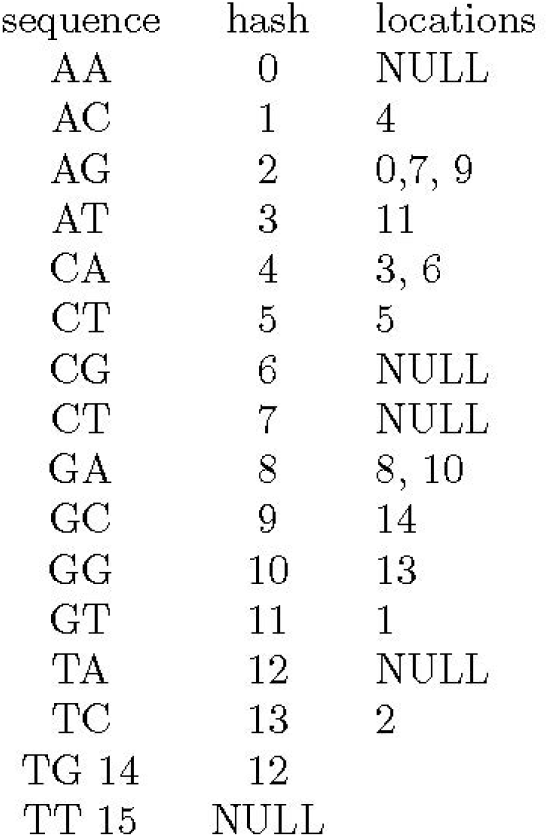
Locations pointed by hash keys for the reference sequence of AGTCACCAGAGATGGC. “NULL” means there is no location for that key.

**Figure 3.**
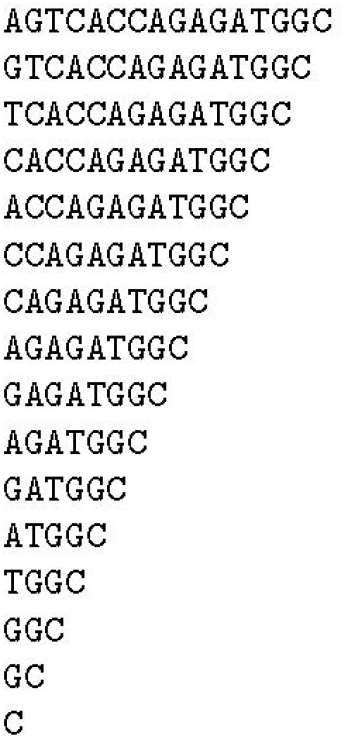
The suffix array before sort for the reference sequence of AGTCACCAGAGATGGC.

If we have sequence AGAGA as a read, for the first AG (at position 0, when we count positions from left and starting with zero), we find candidate locations 0, 7, 9 as shown in figure 2. Here, we shift starting position by one and the key becomes GA, and get 8, 10. Since these locations correspond to position 1 in the read, to obtain locations which correspond to position 0 we should subtract one from these values to obtain 7, 9. For AG at position 2, we have 0, 7, 9 (and thus 5, 7 for the first position of the read), and 8, 10 for GA at position 3 in the read (and thus 5, 7 for the first position of the read). In this case, each key gives multiple values for the first position of the read, but position 7 appears for all four keys and it is the only value shared by all keys. Therefore, we know that AGAGA matches with the reference location 7 and the match is perfect.

If there are SNPs in a read, the hash table gives different results for starting positions which cover the locations of SNPs. If there are inserts/deletions, the hash table gives different (but near) locations for starting positions before inserts/deletions and positions after. Thus, we can get some information on mutations.

This algorithm is quite robust, but the calculation cost per read can become very large. If we make, for example, hash keys of 15 bases, the number of possible values of keys is 4^15^ ≃ 10^9^, and the average number of locations per key is around three since the length of human genome sequence is around 3 × 10^9^. However, some keys appear in a very large number of locations, and that means such keys also appear in many reads. These frequently appearing keys cause a huge increase in the total calculation cost.

For example, if there is one hash key which appears in 10^4^ places in the reference sequence, the probability that one read picks one of these 10^4^ locations is *m* ×10^4^/3 × 10^9^ *m*/3 × 10^5^, where *m* is the length of reads. Thus, if one read has the length of *m* = 150, around one in 2000 reads picks up this pattern and its calculation cost can be 10^4^ times higher than that of other reads. Of course, if there is only one such key, we could just ignore it. In practice, however, there is a spectrum of keys appearing in different number of locations. It is difficult to reduce the total calculation cost using the hash-based algorithm.

### The suffix tree

In principle, if we could use much longer keys, we should be able to avoid this problem of too many matched locations. However, it is unpractical to use the hash length longer than 15, since the amount of memory needed increases exponentially.

One solution for this problem is to use the suffix tree (Weiner, 1973). The suffix tree is a tree structure corresponding to the suffix array, and suffix array is the alphabetically sorted array of all suffixes of the reference sequence. Thus, the suffix array is essentially the array of hash keys with the length the same as that of the reference sequence itself. We can regard the suffix tree as a convenient way to implement very long hash keys.

There are many different ways to make the data structure equivalent to the suffix tree (Farach 1997). Here we present a conceptually simple one for illustration purpose, which is not necessarily practical for actual reference sequence. From the reference sequence of AGTCACCAGAGATGGC, we first make an array of sequences. The first element of the array is the reference itself, and the second element is the same reference but with the first base removed. For *k*-th element we remove first *k* − 1 basess, and thus we have *n* elements, where *n* is the length of the sequence.

Then we sort this array using the dictionary order to obtain the suffix array. The result is shown in figure 4a.

**Figure 4.**
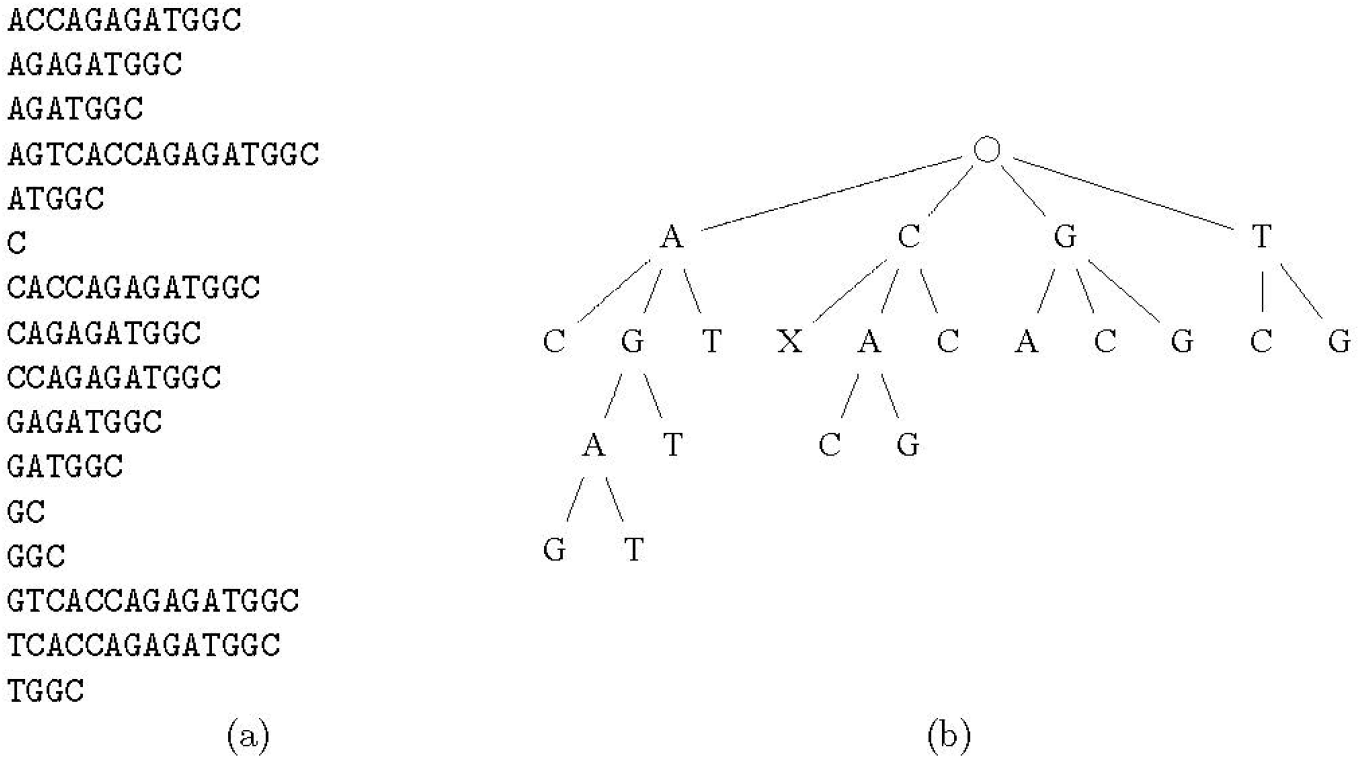
The suffix array (a) and the corresponding suffix tree (b) for the reference sequence of AGTCACCAGAGATGGC. Here “X” in the tree means the end of the sequence.

We can now construct a tree structure corresponding to the suffix array as shown in figure 4b. For one starting location of the read, we can go down the suffix tree to find the location(s) with the longest match. As a result, the number of match locations is dramatically reduced.

Though conceptually simple, the suffix tree has not been widely used for read alignment. One reason is that it requires a large amount of memory. The data size of the reference human genome sequence is around 1GB. The suffix array would need 12 GB or 24 GB (depending on one use 32-bit or 64-bit integer), and the suffix tree can easily consume more than 100GB. Fifteen years ago, when the early versions of widely used genomics programs such as bwa-mem and gatk were designed and developed, the DRAM memory of more than 100GB was very expensive. Moreover, machines which could house a large amount of memory were also very expensive since they had to have a large number of memory slots and thus must use expensive high-end server CPUs and very expensive motherboards.

At that time, it was clearly unpractical to use the suffix tree, since there is an alternative data structure, Burrows-Wheeler Transform (BWT), which is extremely memory efficient. Thus, bwa-mem (Li and Durbin, 2009; Li, 2013, 2012), which is currently the golden standard read aligner, adopted BWT as its basic algorithm and that was where its name, bwa-mem, came from (Burrows-Wheeler Alignment Tool, Maximal Exact Matches). To make use of computers with small amount of memory then available, it was essential to use memory-efficient data structure and the choice to use BWT made perfect sense.

As of 2023, desktop PC motherboards with just four memory slots can house 128GB of memory for less than 1000 USD. Thus, it might be the time to rethink what is the best algorithm for the read alignment. We use the suffix tree itself instead of BWT.

The advantage of the suffix tree is that the algorithm is much simpler compared to the suffix array and BWT, and thus requires a smaller number of the main memory access. To extend the match by one base, the suffix tree algorithm needs to access just one tree node, which is usually a single instance of a class. In contrast, the suffix array and BWT require the accesses to two locations in the suffix array and two more accesses to supporting data structures. Even though the calculation cost itself is not much different, the number of main memory accesses is much smaller for the suffix tree, since the data of one tree node usually fit in the one cache line, while several accesses required by BWT result in the accesses to multiple cache lines. Thus, the extension of the match with the suffix tree is much faster compared to that with BWT, on modern computers with the hierarchical cache structure.

### The hybrid hash-tree algorithm

Even though the suffix tree is quite efficient, it is possible to further improve its efficiency by the following two modifications. The first one is to combine the tree search with the hash key search. We can replace the first *l* levels of the suffix tree with the hash key of the same length, and thus eliminate first *l* − 1 memory access. As we have stated above, *l* =15 is practical with modern computer systems. This modification improves the search speed significantly since, for most cases, the initial hash key search reduces the candidate locations to just a few by single memory access, instead of following the tree structure 15 times. Figure 5 shows the hybrid data structure for the case of *l* = 2.

**Figure 5.**
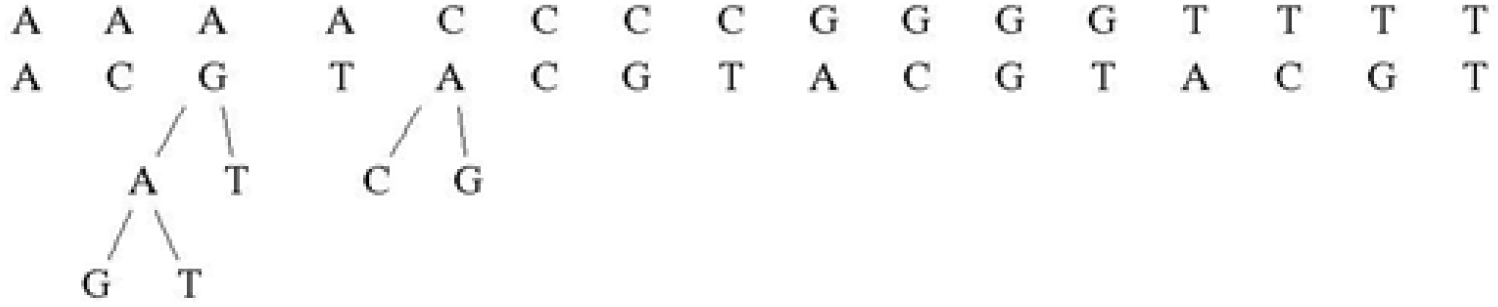
The hybrid hash-tree tree for reference sequence of AGTCACCAGAGATGGC.

Another way to improve the efficiency of the suffix tree algorithm is to collapse multiple levels of tree into a single level. For example, we can replace a two-level tree with four children at each level to one-level tree with 16 children. Using this transformation, we can extend the match by two bases in one iteration, in other words, in one memory access, as far as the single node data fits into one cache line (typically 64 bytes). A tree node with 16 children can fit into 64-byte cache line, while that with 64 children does not. Therefore, instead of a suffix tree with each node corresponding to one base, we use a collapsed tree, with each node corresponding to two bases. This transformation is easy for the suffix tree, but not easy and might be impossible for the suffix array with BWT.

### Search Strategy

With the hash-based algorithm, such as used in BLAST, we obtain location candidates for keys of length l in the read of length m with stride s. Thus, there are (*m* − *l* − *s* + 2)/*s* such keys. With our hybrid algorithm, we could use this same strategy, but it is obviously not ideal. When we find a rather long match, with the next starting position shifted by stride of five or so we will certainly find similarly long match. Also, for the majority of reads the match is ether exact or containing just one SNP. Thus, the calculation cost of the hybrid algorithm can be *O*(*m*^2^). This is certainly not ideal.

We can avoid this problem by simply shift the next starting point by the amount comparable to the match length itself. For example, if we shift the starting position by *p*/2, where *p* is the current match length, the total cost of matching calculation is reduced to *O*(*m*). On the other hand, with this strategy we can miss the longest match, since the true longest match could start at the positions in the read we skipped.

A simple fix for this problem is to retain candidate locations with match length more than half of the apparent maximum match for the score calculation in the next stage. A longer match, if exists, starts at somewhere between the current starting position and the next starting position. In the worst case, where the actual match is the shortest and appears at the leftmost position, it starts at the position next to the current starting position, and extends by two bases after the end position of the current search. Therefore, if we start at the position shifted by *p*/2 from the current starting position, we are can find the latter half of the longest match with length (*p* + 2)/2.

For the search of chimeric alignments, our strategy can be problematic since the candidate regions for a chimeric alignment can be very similar and yet contain, for example, multiple SNPs. Actually, in this case the longest exact match might not give the best matching location either, since the best match location might contain multiple SNPs and the length of the longest exact match location can be short. For such cases, it is necessary to make the shift length small so that we can find all short match locations.

### Extension and scoring

For the extension of the match, we use the usual Smith-Waterman-Gotoh (SWG) algorithm (Gotoh 1982), and we take into account of the Base Quality Score Recalibration procedure (Caetano-Anolles 2023) when assigning the final scores to matches. In our implementation, the actual code for the SWG algorithm uses AVX2 SIMD instruction set, so that we can take advantage of recent processors from both Intel and AMD.

### Parallelization strategy

To make efficient use of modern CPUs, it is essential to make all steps of genome analysis well parallelized for a large number of cores, even when we just use a single desk-side workstation. This is because modern high-end processors have a large number of cores integrated in one package. For example, AMD EPYC 9000 series processors, announced in November 2022, have up to 96 cores in one package, and high-end servers can house two processor packages in one chassis, resulting in 192 cores in one computing node. It is, however, not easy to design a program whose performance scales well for more than 100 cores. Of course, large supercomputers have 1 million or more cores, and at least a few programs can make use of those huge number of cores. However, that is usually for extremely large-scale problems.

From the point of view of parallel processing, one advantage of human whole genome analysis is that there are lots of potential parallelism in all stages of processing. First of all, the alignment of a read can be done independently of those of all other reads. If we use the current typical value of the read length of 150, for a (pair of) fasta files for the read depth of 30 contains 100G bases or around 300M read pairs, and all of these 300M read pairs can be processed in parallel.

There is, however, one practical issue. At least in the case of sample data, fasta files are usually provided as single big text files compressed with gzip. This means that it should be decompressed, and because gzip-compressed file can only be decompressed sequentially, this decompression can take time longer than the rest of the analysis. The actual sequencer should be able to generate many small fasta files for the data of one human genome, since actual reading process is highly parallel. In this paper, we assume that input data are available as a number small fasta files, where the total number of the fasta files is *q*.

Our new hybrid hash-tree algorithm requires fairly large (around 100GB) table to express the reference genome. Therefore, this table must be shared by processes which handle the reads in parallel. We therefore implemented the parallelization using OpenMP. In our implementation, one OpenMP-parallelized loop processes the small fasta files, one pair of files in one iteration. With OpenMP, we can specify how many threads are run in parallel at runtime. Since different threads process different files, there is very low parallelization overhead. Also, we carefully avoided any overhead caused by library calls and memory allocation/deallocation/garbage collection. Thus, as we will see in section 3, we have achieved very good parallel speedup. We are impressed by the fact that Linux operating system can handle huge number of I/O requests quite well. This part of program outputs candidates of matches for each read.

The SWG program, which performs the match through Smith-Waterman-Gotoh algorithm and calculates the matching sore, does not require large tables. Therefore, we implement this part as a single-thread program, which process one file and generate SAM-format output. For parallel processing we just run a fixed number of this program in parallel.

The output of the SWG program is then processed by the program for variant calling. The parallelism is also there for variant calling, but it is not of the level of reads but of the level of the locations in the reference genome. Conceptually, what we do for variant calling is, for each base location in the reference genome, to see if the bases of reads aligned at its location contains SNP or other mutations. To do this, we need to be able to find all reads which cover the location we are looking at, and this is most easily done by sorting reads according to the aligned locations.

Usually sorting of reads is done using samtools. Here, again, the processing time of samtools sort can be longer than the rest of processing, and it is important to speed up the sorting process. We have implemented parallel off-the-core bucket sort. We divide the reference locations into *r* regions (*r* ∼ 1000 or more). The SWG program creates *r* output files, and writes each SAM record to the file with the appropriate region. Since there are *q* small fasta files, we will have *pq* small SAM files. If we call one SAM file generated from fasta file *i* for reference region *j* as *S*_*ij*_, we can obtain all SAM records for region *j* by combining all *S*_*ij*_ for 0 ≤ *i* < *q* - 1. The variant caller reads these files and sorts SAM record on memory.

### Variant Calling

As we stated in the previous subsection, our variant caller accepts the SAM records in a specified range of the reference sequence and performs sorting. Then it performs so called MarkDuplicates, and checks if there is SNP or INDEL for each location in the reference region.

### Hardware and Software Platform

For all test calculations in this paper we used a Linux server with AMD ThreadRipper 3990X 64-core processor, 256GB of DDR4 main memory, and 4TB of PCIe gen3 M.2. SSDs.

The operating system is Ubuntu 20.4 LTS. We used gcc version 9.4.0 (Ubuntu 9.4.0-1ubuntu1 20.04.1) which comes with the operating system.

## Results

### Benchmark problem

As the benchmark problem, we used the data provided for PrecisionFDA Truth Challenge V2 (Olson et al., 2022). We follow the challenge contest procedure and show the accuracy and timing result. In appendix, we show the command parameters used for bwa-mem/gatk processing.

### Accuracy

Table 1 give the accuracy result, as was required in the PrecisionFDA Truth Challenge V2 contest. For each of three sample data, HG002, HG003 and HG004, we present the result with bwa-mem/gatk, as well as two results with our system. One is with the high-accuracy mode, and the other with the low-accuracy, fast mode. As in the case of the original Truth Challenge V2 contest, some of the adjustable parameters for our system was tuned to give the best result for HG002, and the same set of parameters are used for HG003 and HG004. We can see that the final F-measure values of our high-accuracy results are slightly better than that of bwa-mem/gatk, while those of low-accuracy result are slightly worse. The difference is, in all cases, quite small, and we can safely conclude that our system has achieved the accuracy of the same level as that of bwa-mem/gatk.

**Table 1.** Accuracy test results.

| (a): HG002 |  |  |  |  |  |  |  |
| --- | --- | --- | --- | --- | --- | --- | --- |
|  | True positives | True calls | False-pos | False-neg | Precision | Sensitivity | F-measure |
| Gatk | 3854404 | 3855258 | 54542 | 37091 | 0.9860 | 0.9905 | 0.9883 |
| High | 3843485 | 3845450 | 38814 | 48010 | 0.9900 | 0.9877 | 0.9888 |
| Low | 3840691 | 3841762 | 44070 | 50804 | 0.9887 | 0.9869 | 0.9878 |
| (b): HG003 |  |  |  |  |  |  |  |
|  | True positives | True calls | False-pos | False-neg | Precision | Sensitivity | F-measure |
| Gatk | 3795148 | 3795094 | 53773 | 36846 | 0.9860 | 0.9904 | 0.9882 |
| High | 3785845 | 3786914 | 36724 | 46149 | 0.9904 | 0.9880 | 0.9892 |
| Low | 3783096 | 3783268 | 41959 | 48898 | 0.9890 | 0.9872 | 0.9881 |
| (c): HG004 |  |  |  |  |  |  |  |
|  | True positives | True calls | False-pos | False-neg | Precision | Sensitivity | F-measure |
| Gatk | 3819599 | 3819528 | 54716 | 37529 | 0.9859 | 0.9903 | 0.9881 |
| High | 3808201 | 3809240 | 39099 | 48927 | 0.9898 | 0.9873 | 0.9886 |
| Low | 3805555 | 3805753 | 43751 | 51573 | 0.9886 | 0.9866 | 0.9876 |

Table 2 shows the accuracy for SNPs and INDELs (non-snps), for the case of HG002 sample. Our result is slightly better than that of bwa-mem/gatk for SNPs and slightly worse for INDELs. Again, the difference is very small.

**Table 2.** HG002 SNP/INDEL result.

|  | true positives | false positives | true positives calls | false negatives | precision | sensitivity | f-measure |
| --- | --- | --- | --- | --- | --- | --- | --- |
| Gatk SNP | 3336055 | 45856 | 3337490 | 28373 | 0.9864 | 0.9916 | 0.9890 |
| Gatk INDEL | 518349 | 8686 | 517768 | 8718 | 0.9835 | 0.9835 | 0.9835 |
| High SNP | 3325657 | 29106 | 3329961 | 38771 | 0.9913 | 0.9885 | 0.9899 |
| High INDEL | 517828 | 9708 | 515489 | 9239 | 0.9815 | 0.9825 | 0.9820 |

Compared to other results presented in Olson et al. (2022), our results (and that of bwa-mem/gatk) are of course not quite the best, even within non-deep-learning results, but are close to the average of them, since the result for Illumina data ranges from 97% to 99.7%.

### Timing

Tables 3 and 4 show the elapsed time to process HG002 sample using our system and bwa-mem/gatk, respectively. For both cases, the time for each command and the total time are shown. For our system, times for both the high-accuracy and low-accuracy modes are shown. When we compare the total time, our low-accuracy mode is 49.5 times faster than bwa-mem/gatk, while our high-accuracy mode is 26.4 times faster. As we have stated earlier. bwa-mem uses less than 10% of the total time. If we compare the time for bwa-mem alone and that for our aas-match part, which does roughly what bwa-mem does, our low-accuracy mode is 4.4 times faster and high-accuracy mode is 2.2 times faster than bwa-mem, respectively.

**Table 3.** Time in minutes to process HG002 sample using our system.

|  | High accuracy | Low accuracy |
| --- | --- | --- |
| aas-match | 55.82 | 28.32 |
| aas-vcall | 2.68 | 2.92 |
| Total | 58.50 | 31.23 |

**Table 4.** Time in minutes to process HG002 sample using bwa-mem/gatk.

|  |  |
| --- | --- |
| bwa-mem | 124.40 |
| MarkDuplicates | 135.75 |
| BaseRecalibrator | 203.67 |
| PrintReads | 716.67 |
| HaplotypeCaller | 363.83 |
| Total | 1544.32 |

For the procedures after bwa-mem, with our system everything is done in a single command, aas-vcall, in less than three minutes. It effectively does the sorting of SAM records, MarkDuplicates, BaseRecalibration, and variant calling. Compared to gatk tools which take around 24 hours in total, our system is around 500 times faster. If we compare the time for HaplotypeCaller only, our system is still 120 times faster. This difference is not due to any new algorithm, but primarily because everything is written in C language in such a way that there is no serious parallelization overhead.

Table 5 show the throughput of systems in terms of TBp/day, for present work, NVIDIA V100 (Franke and Crowgey, 2020) and Fujitsu A64fx (Suzuki et al., 2021). We can see that our system is by far more cost-effective compared to systems reported in recent works. With our software, a single-processor workstation achieves the throughput comparable to (but faster than) those of 16 NVIDIA V100 GPGPUs or 96 Fujitsu A64fx processors.

**Table 5.**
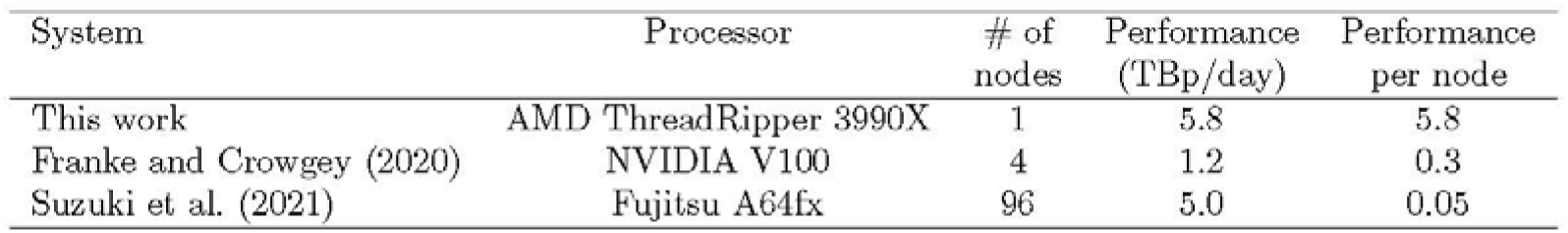
Throughput of systems in terms of TBp/day.

| System | Processor | # of nodes | Performance (TBp/day) | Performance per node |
| --- | --- | --- | --- | --- |
| This work | AMD ThreadRipper 3990X | 1 | 5.8 | 5.8 |
| Franke and Crowgey (2020) | NVIDIA V100 | 4 | 1.2 | 0.3 |
| Suzuki et al. (2021) | Fujitsu A64fx | 96 | 5.0 | 0.05 |

### Parallel Efficiency

Figure 6 shows the time to process HG002 sample data as the function of the number of threads used. Dashed line indicates the ideal parallel speedup. We can see that processing speed is almost proportional to the number of threads, for up to 64 threads. For more than 64 threads, the gain is small since the physical number of cores of the processor we used is 64.

**Figure 6.**
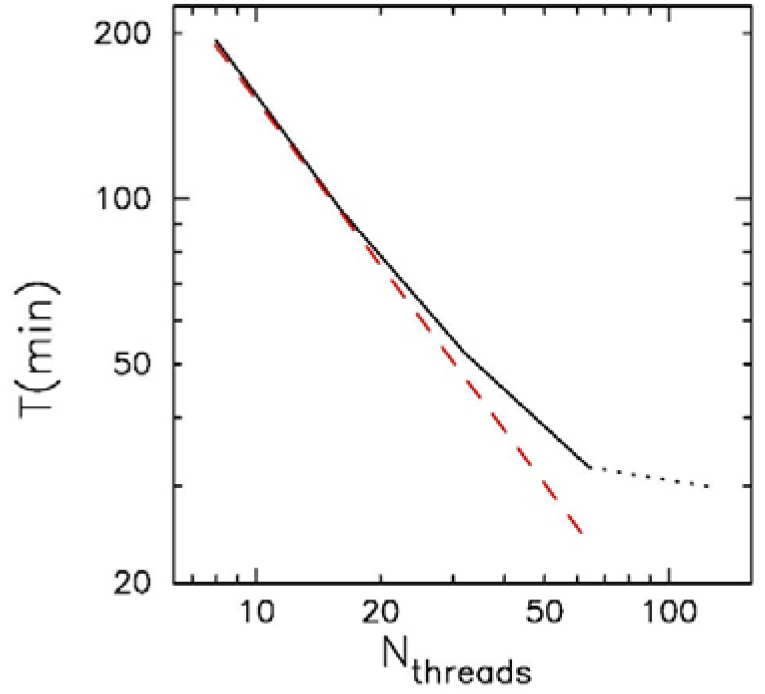
Time in second T to process HG002 sample data as the function of the number of threads. Dashed line indidate ideal scaling. For, the result is shown in dotted curve since the physical number of cores is 64.

The main reason why the speedup is somewhat less than ideal for large number of cores is because of the limitation of the clock frequency. On AMD ThreadRipper 3990X processor, when only a small number of cores are active, their clock frequency can reach to 4.3 GHz. However, when a large number of cores are used, the clock frequency goes down due to the limit in the power consumption. We found that the clock frequency was 3.9GHz for 32 threads, and 3.3GHz for 64 and 126 threads. This reduced clock frequency explains why the parallel speedup is not ideal.

## Discussions

In this paper, we present the algorithm, implementation and performance of our fast aligner for short reads. Our system is two to four times faster than bwa-mem on the same computer system. The accuracy achieved for dataset used in Precision FDA Truth Challenge V2 is very close to that of bwa-mem/gatk for our low-accuracy mode (F-measure difference of 0.01-0.05%), and slightly more accurate for our high-accuracy mode.

We also developed a new implementation of the variant caller, which is used to obtain the above result. It can process Truth Challenge V2 samples in less than three minutes on a 64-core AMD ThreadRipper processor. Our variant caller is more than 100 times faster than gatk Haplotype-Caller.

Right now, our system has been tested only for germline mutations and not well tested for somatic mutations or structural variants. We are currently working on the efficient detection of structural variants. Here, bwa-mem does consume a significant fraction of the total processing time. Even so, the total speed-up factor is rather limited, since variant callers for structural variants consume time comparable to that consumed by bwa-mem. Thus, to improve the total performance, it is necessary to improve the performance of the variant caller here as well. There are still many rooms of performance improvement in our new aligner, and we hope to report on improved performance in near future.

## Acknowledgments

This work was done under the research contract between Advanced Accelerating Systems Co. Ltd. and K. K. Dnaform.

## Authors’ contribution

Conceptualization: JM, TE, RH, YH

Data curation: JM

Formal analysis: JM

Funding acquisition: YH, RH

Methodology: JM

Writing – original draft: JM

Writing – review & editing: JM, TE

**Appendix A: Command Scripts**

Command Scripts for our system Listing 1: Commands for our system

**Listing 1:**
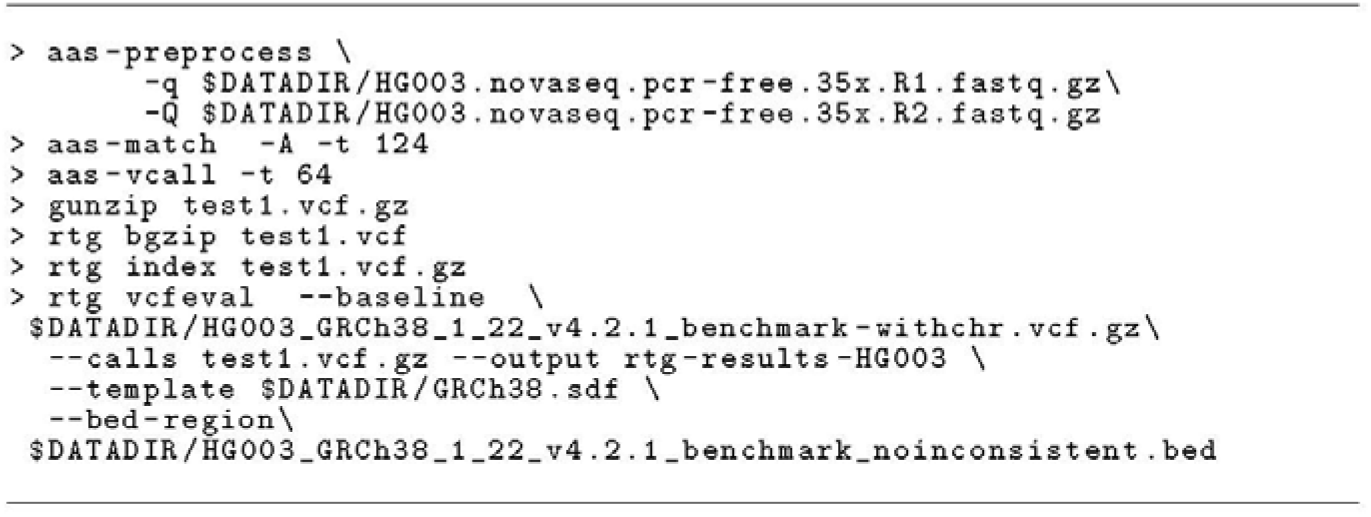
Commmands for our system

Command Scripts for bwa-mem/gatk Listing 2: Commands for bwa-mem/gatk>

**Listing 2:**
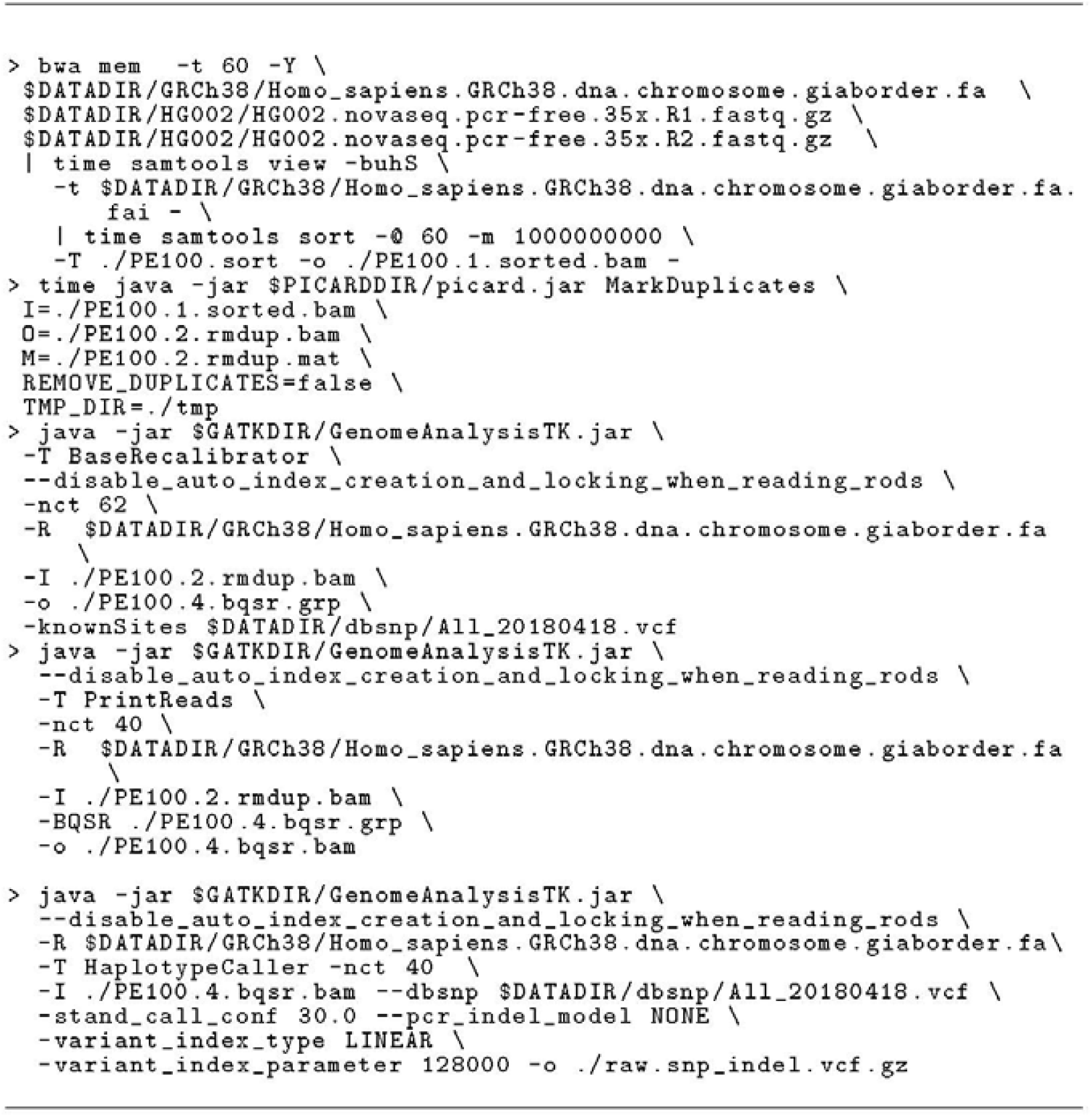
Commmands for `bwa-mem/gatk`

